# The R-loop Grammar predicts R-loop formation under different topological constraints

**DOI:** 10.1101/2024.12.03.626533

**Authors:** Margherita Maria Ferrari, Svetlana Poznanović, Manda Riehl, Jacob Lusk, Stella Hartono, Georgina González, Frédéric Chédin, Mariel Vázquez, Nataša Jonoska

**Affiliations:** University of Manitoba, Winnipeg, MB CA; Clemson University, Clemson, SC USA; Rose-Hulman Institute of Technology, Terre Haute, IN USA; University of California, Davis, Davis, CA USA; University of South Florida, Tampa, FL USA

**Keywords:** R-loops, Formal Grammars, DNA topology, Transcription

## Abstract

R-loops are transient three-stranded nucleic acids that form during transcription when the nascent RNA hybridizes with the template DNA, freeing the DNA non-template strand. There is growing evidence that R-loops play important roles in physiological processes such as control of gene expression, and that they contribute to chromosomal instability and disease. It is known that R-loop formation is influenced by both the sequence and the topology of the DNA substrate, but many questions remain about how R-loops form and the 3-dimensional structures that they adopt. Here we represent an R-loop as a word in a formal grammar called the *R-loop grammar* and predict R-loop formation. We train the R-loop grammar on experimental data obtained by single-molecule R-loop footprinting and sequencing (SMRF-seq). Despite not containing explicit topological information, the R-loop grammar accurately predicts R-loop formation on plasmids with varying starting topologies and outperforms previous methods in R-loop prediction.

**Author summary:** R-loops are prevalent triple helices that play regulatory roles in gene expression and are involved in various diseases. Our work improves the understanding of the relationship between the nucleotide sequence and DNA topology in R-loop formation. We use a mathematical approach from formal language theory to define an R-loop language and a set of rules to model R-loops as words in that language. We train the resulting R-loop grammar on experimental data of co-transcriptional R-loops formed on different DNA plasmids of varying topology. The model accurately predicts R-loop formation and outperforms prior methods. The R-loop grammar distills the effect of topology versus sequence, thus advancing our understanding of R-loop structure and formation.

## Introduction

R-loops are three-stranded structures composed of an RNA:DNA duplex and a single-strand of DNA. Initially discovered in bacteria, R-loops constitute 3-5% of the genome of yeasts, plants, and mammals [1, 2, 3, 4, 5, 6] and are at least one order of magnitude longer than other non-B DNA multi-stranded nucleic acid structures [7, 8]. R-loops form co-transcriptionally when the nascent RNA invades the DNA duplex and the RNA hybridizes with the template DNA strand [9]. The unpaired non-template DNA strand is free to wrap around the hybrid duplex or to fold upon itself into a secondary structure (Figure 1). R-loops arise through a dynamic process that begins with DNA duplex invasion by the nascent RNA behind the advancing RNA polymerase (*initiation phase*). Once an R-loop has been seeded, it can extend dynamically during transcription (*elongation phase*). Having reached a point at which the structure can no longer grow, the R-loop terminates (*termination phase*). Termination may be followed by an equilibration process, where the exact boundaries of the structure may shift through branch migration [10]. Eventually, the R-loop dissociates and the B-form DNA duplex is restored. Figure 2 shows the different stages of R-loop formation.

**Figure 1.**
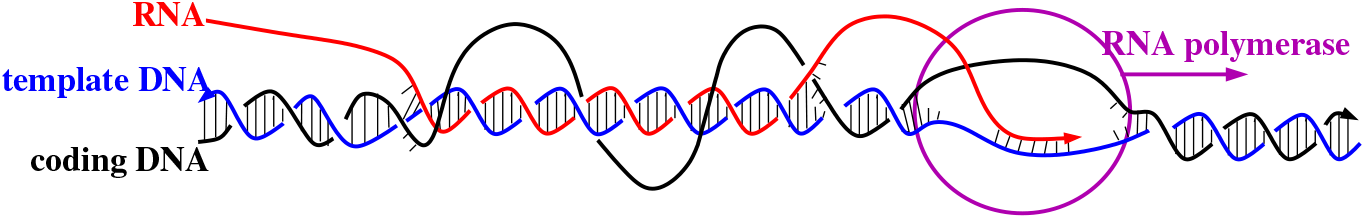
Co-transcriptional R-loops. Transcription of DNA into RNA is mediated by the RNA polymerase. A co-transcriptional R-loop forms behind the polymerase when the RNA transcript invades the double-stranded DNA (dsDNA) and hybridizes with the template DNA strand. The template and non-template DNA strands are shown in blue and black, respectively. The red strand represents the RNA transcript. In the R-loop, the non-template DNA strand is unpaired and free to wrap around the RNA:DNA duplex. The 3′-ends are indicated by an arrowhead.

**Figure 2.**
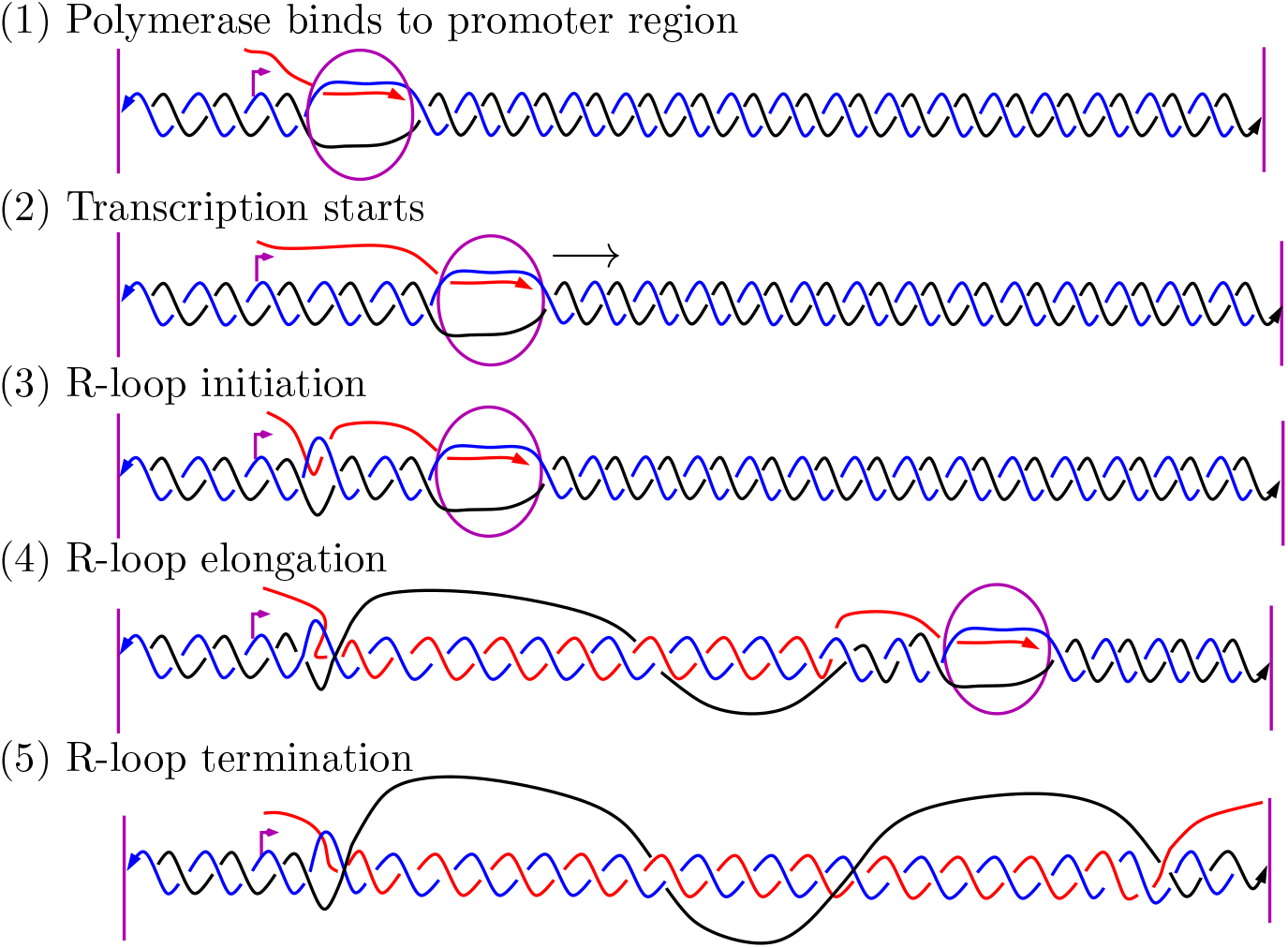
Stages of R-loop formation. (1) The polymerase binds to the promoter sequence (purple arrow). (2) Transcription starts. The polymerase moves from left to right and generates the RNA transcript (in red) in the 5′ to 3′ direction. (3) R-loop initiation: the nascent RNA invades the dsDNA and hybridizes to the DNA template strand (in blue). (4-5) R-loop elongation and eventually termination.

Organisms have evolved complex pathways that regulate R-loop levels [11]. Genome mapping studies indicate that R-loops do not form randomly [12, 9]. DNA sequence analysis, biochemical experiments and statistical mechanical modeling suggest that both the DNA sequence and the topology play key roles in promoting and controlling R-loop formation [9, 13]. The fundamental forces that drive R-loop initiation, elongation and termination remain under-studied.

In formal language theory, a grammar is a set of production rules that generate strings in a formal language. Applications of formal grammars can be found in a wide range of areas such as theoretical computer science, theoretical linguistics, and molecular biology. In molecular biology, applications include modeling regulation of gene expression [14], gene structure prediction [15], sequence analysis [16] and RNA secondary structure prediction [17].

Here we introduce the *R-loop grammar*, a predictive formal grammar model of R-loop formation that advances our understanding of the structure, formation and biological function of R-loops. We train and test the R-loop grammar on single-molecule RNA footprinting and sequencing (SMRF-seq)[18, 8] experimental data to define the syntax of an R-loop language. Each R-loop is written as a word in this language. The grammar model predicts the probability that an R-loop will form in a given DNA segment. It also predicts the location and basic 3-dimensional (3D) structure of the R-loop (e.g. coiling of the single DNA strand around the duplex).

We study R-loops generated using SMRF-seq upon *in vitro* transcription of two plasmids, pFC53 and pFC8 [13, 18, 8]. This method is unbiased, ultra-high coverage, strand-specific and yields high-resolution results. Unlike other methods (e.g. DRIP) that output population averages, SMRF-seq on *in vitro* transcribed DNA templates provides information about R-loops at nucleotide resolution. The experimental data include information on R-loop formation under three topological conditions: linear, negatively supercoiled and hyper-negatively supercoiled.

For each plasmid and each starting topology, we train the R-loop grammar on a portion of the data. We use the experimental data to generate the syntax of the new language and obtain probabilities for the production rules. Figure 3 shows how the substrate topology of pFC53 influences the formation of R-loops. The corresponding figure for pFC8 is Figure S2 in the Supporting Information (SI) Appendix. R-loops cluster in two regions for all substrate topologies. However, R-loop initiation shifts to the left as the supercoiling levels increase. As observed in [13], the majority of the R-loops in the hyper-negatively supercoiled plasmid appear closer to the transcription start; this is not the case for the other two topologies (Figure 3).

**Figure 3.**
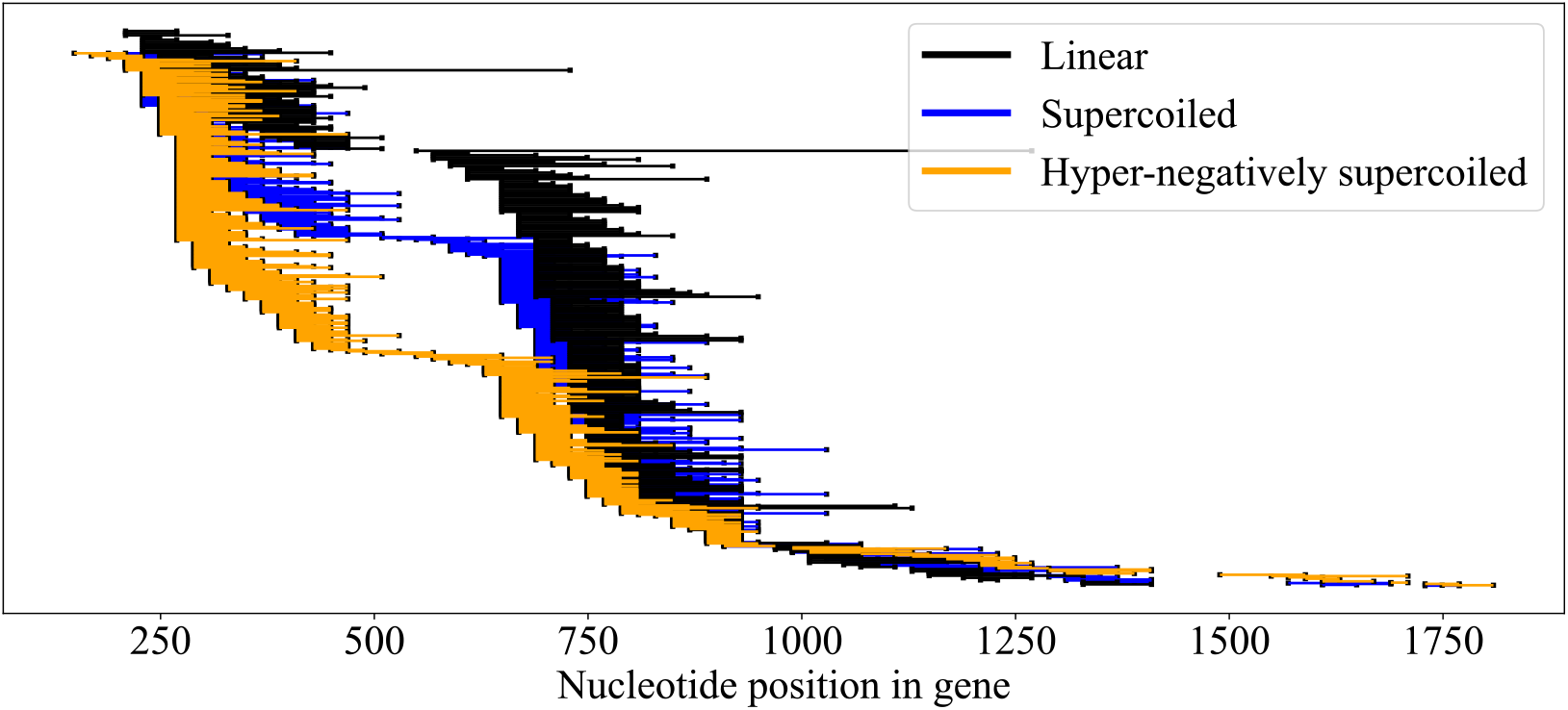
Experimental R-loop locations for plasmid pFC53 with starting topology: linear (black); supercoiled (blue); and hyper-negatively supercoiled (orange). The x-axis indicates the nucleotide position of the gene starting at 0 rounded to the nearest 20th nucleotide. Each horizontal line segment corresponds to one experimentally detected R-loop. The R-loops have been sorted by the starting nucleotide. Each data set is uniformly spread vertically (255 R-loops for linear, 612 for supercoiled and 408 for hyper-negatively supercoiled). In this way proportional differences in R-loop initiation under the three conditions can be observed independent of the number of experimental R-loops observed.

We show that the R-loop grammar model distills the effect of DNA topology on R-loop formation. Unlike the R-looper prediction method [13], the model does not explicitly include parameters to specify the topology of the substrate. Instead, the model learns about these topological constraints from the data. The key findings are that the substrate topology affects the probability of the production rules and consequently the R-loop grammar effectively predicts the probability of R-loop formation.

### The R-loop Grammar: a Model for R-loop Formation

#### Formal grammars and R-loops

A formal grammar consists of a finite set of symbols partitioned into *variables V* and *terminals* Σ, and a finite set of *production rules* {*u* → *v*}. When applying the rule *u* → *v* on a word *xuy*, the subword *u* is substituted by the subword *v* yielding a word *xvy*. A word derived by the grammar is obtained by a consecutive application of rules starting from *S*, a non-terminal symbol designated as a *starting symbol*. The language generated by the grammar consists of all words comprised of terminal symbols that can be derived by the rules starting from *S* [19, 20, 21].

We define the *R-loop grammar* as a formal grammar whose terminal symbols correspond to the basic structures of an R-loop. The symbol *α* describes the R-loop initiation (RNA invasion), *ω* describes the termination (RNA dissociation). The symbols *σ* and 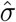 represent short DNA:DNA hybrids with a free RNA strand (the RNA transcript). The symbols *τ* and 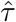 represent RNA:DNA hybrids with a free DNA strand (the non-template strand). The ‘^’ indicates a more stable configuration, i.e. a configuration that is not prone to changing state. Therefore 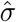 denotes a structure unlikely to transition from a DNA duplex to an RNA:DNA hybrid, and 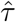 denotes a structure unlikely to transition from an RNA:DNA hybrid back into the DNA duplex. Figure 4 shows the main terminal symbols in the R-loop grammar. Figure 5 shows a word generated by an R-loop grammar and its corresponding R-loop structure. Note that if the sequence stability weakens within an R-loop, a less stable RNA:DNA duplex (indicated by *τ*) may follow after an initial string of one or more 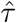’s. Intuitively, one or more consecutive *τ* ‘s may lead to an R-loop termination region. The three-strand sections *α* and *ω* correspond to those regions of branch migration that mark the initiation and termination of the R-loop, respectively.

**Figure 4.**
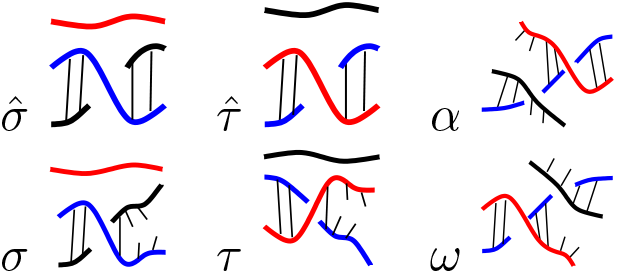
Basic 3-strand structures found in an R-loop and their associated symbols in the R-loop grammar. We indicate less stable configurations *σ* and *τ* by breakage in the hydrogen bonds. This representation should not be taken as literal breakage of all bonds in that vicinity, but rather as an indication that this region is unstable and prone to opening of the helix. The color coding is as in Figure 1.

**Figure 5.**
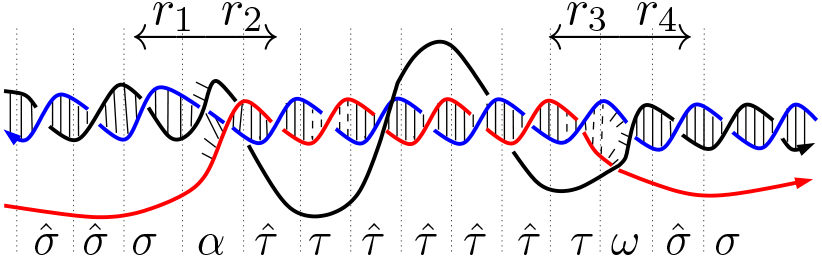
An R-loop associated with the word 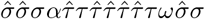. The colors are as in Figure 1. For simplicity, we omit the broken hydrogen bonds for *σ* and *τ* and simply indicate them under the diagram.

We use a training set extracted directly from the experimental data to determine the assignment of terminal symbols in the R-loop grammar and the application of each production rule. The application of each production rule depends on a probability distribution generated from the data. We apply this approach to SMRF-seq data for plasmids pFC53 and pFC8 with three different plasmid topologies.

#### Symbol assignment and R-loop production rules

We use the experimental data first reported in [13] to determine the probabilities for the production rules. To each block of *k* consecutive nucleotides in the sequence (*k*-mer) we assign a terminal symbol according to the probability that it is contained in an R-loop. The words generated by the grammar correspond to R-loops (Figure 5). All the experimental data is included on our GitHub [22].

Occasionally the basic terminal symbols cannot be unequivocally assigned to a *k*-mer based on the available experimental data. In those instances we expand the set of terminal symbols to accommodate the corresponding *k*-mers. The symbol *δ* (respectively, *β*) represents ambiguous *k*-mers that, according to the statistical analysis of the training data, could be associated with both *σ* and 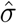 (respectively, *τ* and 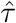). The *k*-mers outside (respectively, within) an R-loop for which the statistical analysis does not provide enough information are indicated with *γ* (respectively, *ρ*). Moreover, to account for the fact that the experimental assignment of initiation and termination of each R-loop is not precise [8], we substitute the terminal *α* (resp. *ω*_*i*_) with a sequence of terminal symbols *α*_*i*_ (resp.,*ω*_*i*_) corresponding to an initiation (resp., termination) site of length *i*, where *i* = 0,…, *k*−1.

As is common in formal grammars, the non-terminal symbols are written with capital symbols, while rules *X* → *Y* and *X* → *Z* with the same left side are written as *X* → *Y* | *Z*. For *i* = 0,…, *k* −1 we define the following rules:

1. start rule

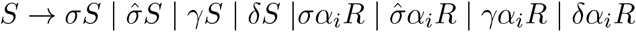
2. RNA:DNA duplex

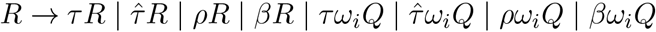
3. DNA:DNA duplex

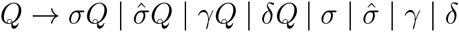

The sequence analysis in the Materials and Methods section describes a way to map each k-mer to a terminal symbol. With such an assignment, each R-loop can be represented as a word over the symbols in this grammar. The word can be obtained by starting with the symbol *S* and applying the rules above in a unique way. For example, by applying the rules 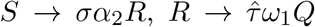, and *Q* → *σ* in succession, one obtains the word 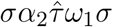. We explain the probability assignments for each production rule in the Materials and Methods section and in the SI Appendix 3. This further associates a probability to each R-loop occurrence that is computed as the product of the probabilities of the corresponding production rules. For example, the probability of the R-loop described by the word 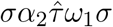 is

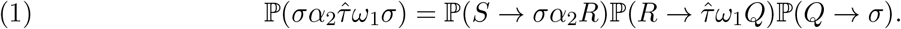

#### General approach

One goal when analyzing the experimental R-loop data obtained from the plasmids pFC8 and pFC53 is to identify and clarify DNA sequence patterns specific to the initiation, elongation, and termination of R-loops. For each R-loop in the training set, we identify four regions of interest, *r*_1_ and *r*_2_ immediately upstream and downstream of the R-loop initiation, as well as *r*_3_ and *r*_4_ immediately upstream and downstream of the R-loop termination (Figure 5). In each region *r*_*i*_ we consider all of the *k*-mers that appear in that region and we assign weights according to the relative frequency of the respective *k*-mer (see Materials and Methods). We form a dictionary for each region by using these weights to associate a terminal grammar symbol to each k-mer. The single plasmid dictionaries showed a significant proportion of *k*-mers as indeterminate symbols (e.g. *γ, ρ*) ranging from 15.0% to 30.2%. To improve the prediction we incorporate the information from both plasmids by taking a union of their dictionaries. With this approach, the proportion of indeterminate symbols in the union of dictionaries lowered to ranges from 4.8% to 13.4% (find details in SI Appendix Table S1). When taking the union, we consider two ways of resolving conflicting information in symbol assignment, stochastic (coin toss) and deterministic, guided by the description of the grammar symbols (see SI Appendix 2).

For each plasmid and each starting topology, we take a subset (10% of 2*β*) of the data as a basis for training the grammar (see Results as well as Materials and Methods). We translate the R-loop nucleotide sequences from the training set as words over the grammar symbols and reverse-engineer the sequences of production rules that generate the words. We assign probabilities to the rules according to the frequencies of each rule application. To make predictions, we first use the grammar to generate all possible R-loop words for a given plasmid. The probability 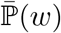 of a word *w* is proportional to the product of the probabilities of the sequence of rules that generate *w* (e.g. (1)). We compute the probability of each nucleotide being in an R-loop by summing the probabilities of the words where this event occurs. For example, let *q*_*i*_(*w*) be 1 if the *i*-th nucleotide is in the R-loop represented by *w*, and 0 otherwise. The probability 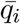 that the *i*-th nucleotide is in an R-loop is

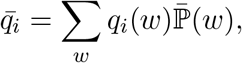

where the sum is taken over all possible R-loop words in a dataset. Finally, we report the average probability that each nucleotide is in an R-loop (see Results and Materials and Methods).

## Results

We analyzed three datasets for each plasmid that differ in their topology before transcription: *linear*, where the plasmids are linearized before transcription; *supercoiled*, where the plasmids have the native supercoiling from bacteria, i.e. a supercoiling density of ∼-0.07; and *hyper-negatively supercoiled*, where the plasmids are treated with gyrase before transcription to double the super-coiling density to ∼ − 0.14 [23].

We generated predictions using R-loop grammars obtained from a stochastic union of dictionaries and from a deterministic union. In this section we discuss results from the stochastic method (Figures 6 and 7). The predictions with the deterministic method showed negligible differences (SI Appendix Figure S5). Figure S4 shows stochastic assignments for the full set.

**Figure 6.**
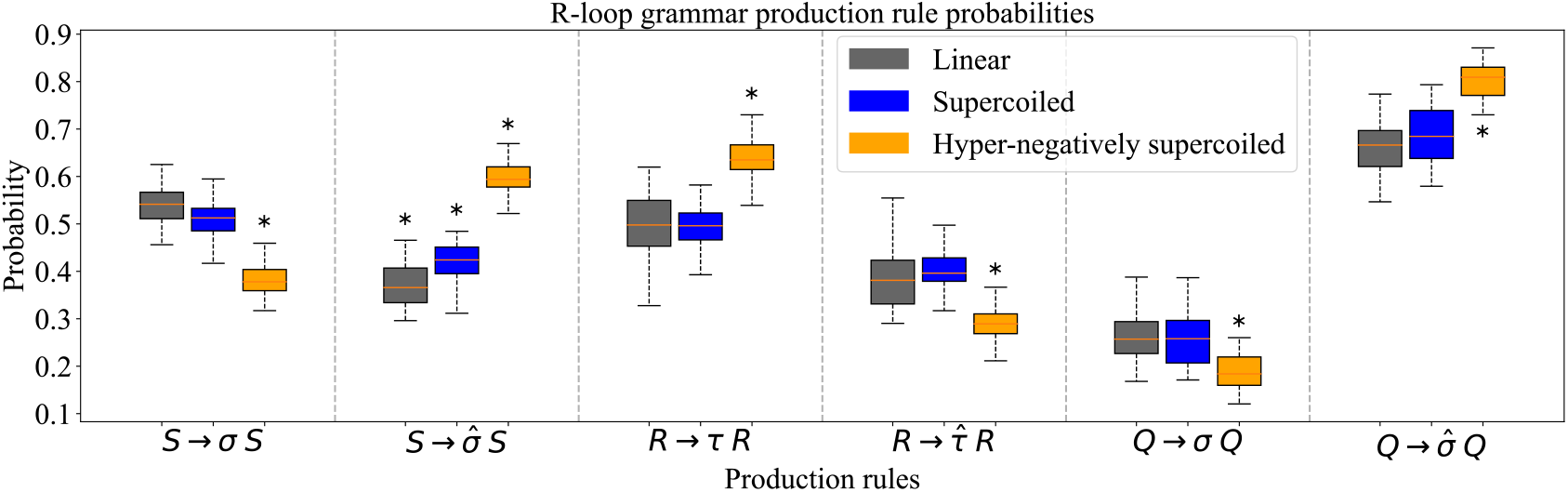
Production rule probabilities. The boxplots illustrate the changes in the probabilities for the six main production rules related to the stability of the structure before, within, and after an R-loop as the topology from the substrate changes from linear to hyper-negatively supercoiled. The probabilities are obtained for a grammar defined for the union of dictionaries with parameters *k* = 4 and *p* = 13. The mid-line of each box is the median, with the first and third quartiles indicated by the box frames. The whiskers represent the largest point not more than 1.5 interquartile range (IQR) beyond the box frame. An asterisk *indicates that the difference is significant against the results from the other two topologies (p ≤ 0.006).The significance of these probability changes are obtained with Bonferroni adjusted p-values ≤ 0.006 according to the pairwise T-test. (See Table S3 for precise values.)

**Figure 7.**
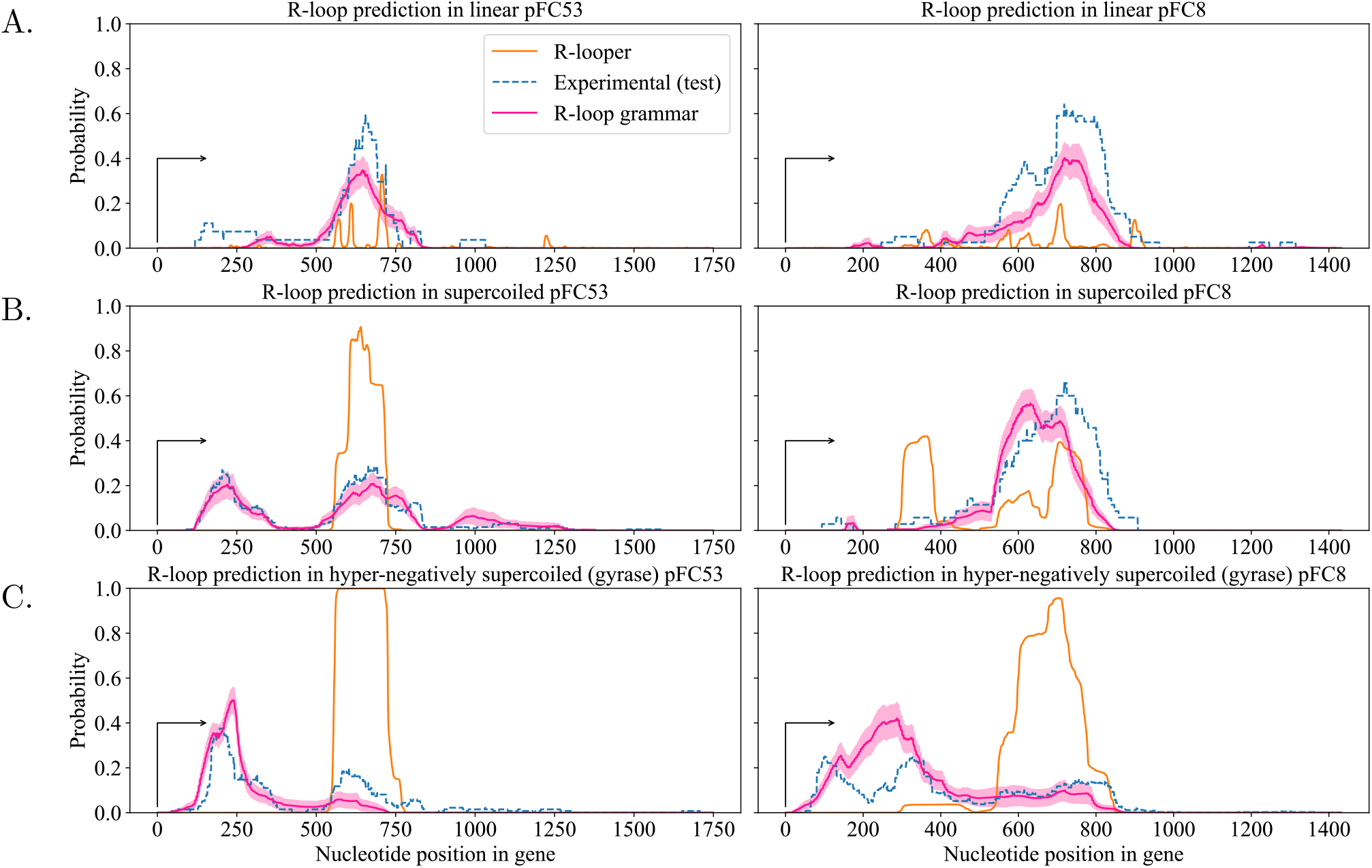
Experimental data (holdout set, dashed blue), R-loop grammar model (pink) and R-looper probabilities (orange), for different topologies on plasmids pFC8 and pFC53. For each plasmid, the figure shows the R-loop grammar predictions obtained with the union of dictionaries for *k* = 4,*p* = 13, as an ensemble average of 30 models. The pink shaded area represents the standard error of the mean. The experimental data show the number of R-loops that contain a particular nucleotide within an R-loop, divided by the total number of R-loops in the holdout set. The arrow indicates the start of transcription; nucleotides are enumerated from that position on the *x*-axis. The substrate topology is indicated in each graph: linear (top row); supercoiled (middle row); hyper-negatively supercoiled (bottom row).

### Plasmid topology drives the probability of the production rules

Figure 6 shows the probability assignments for the production rules associated with symbols 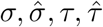 across all datasets. The probabilities of the other productions rules are included in SI Appendix Figure S3a. A higher probability for a given rule implies that the training set contains a larger number of *k*-mers associated with that symbol. If a *k*-mer repeats, its multiplicity is taken into account. Note that the production rule probabilities change significantly with plasmid topology, which is consistent with the premise in [13].

As the supercoiling level increases towards hyper-negatively supercoiled, the probability of a stable DNA duplex 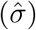 outside the R-loop also increases (rules 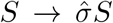 and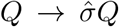). This suggests that the *k*-mers outside the R-loop region are well determined. Once an R-loop starts, the pattern is reversed and the probability of elongating a stable R-loop decreases as supercoiling levels increase (rule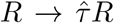). We validated these observations using Kendall’s Tau correlation coefficient (SI Appendix Table S4). Note that in comparison with linear and supercoiled plasmids, a significantly higher number of R-loops in hyper-negatively supercoiled plasmids appear closer to the transcription start (Figure 3). Because our dictionary assignment is focused on the regions at the start and the end of the R-loops, in hyper-negatively supercoiled plasmids a larger number of *k*-mers are spread throughout the R-loop containing regions (e.g. region between ∼ 250nt and ∼1250nt for pFC53, Figure 3). This spread of the R-loop starting points in hyper-negatively supercoiled cases implies that *k*-mers within those R-loops have a somewhat weaker association with R-loops. Hence, rule *R* → *τR* occurs with a much higher probability than 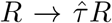. If we instead focus on linear and supercoiled plasmids, we observe that k-mers are mostly concentrated around the peak of the R-loop clusters (∼ 650nt to ∼ 1250nt). Accordingly, the difference between probabilities *R* →*τR* and 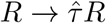 is much smaller.

After the R-loop terminates, the probability of transitioning to a stable DNA duplex is highest for all topologies. We observe the same trends in probabilities upon training the grammar on the data from each plasmid separately (SI Appendix, Figures S3b and S3c), as well as for the deterministic assignment of the symbols (SI Appendix, Figure S3d).

### The R-loop grammar accurately predicts R-loop formation for different topologies

The R-loop grammar model has two adjustable parameters, the tuple size *k* and the padding length *p*. Due to experimental sensitivity, the initiation of the R-loop may vary up to 15 nucleotides from the location observed through SMRF-seq [8]. To account for this, we focused on padding parameters *p* = 7, 13. The k-mer plus the padding correspond approximately to one (*p* = 7), or one and a half (*p* = 13) turns of an A-DNA double-helix (∼11bp). We start with a premise that the k-mers in the vicinity of the experimental R-loop start/end locations are critical for accurate prediction. We add the padding (nucleotide segments) before and after the R-loop start/end, thus defining the k-mers in regions *r*_1_ to *r*_4_ (see Figure 8 in Materials and Methods).

**Figure 8.**
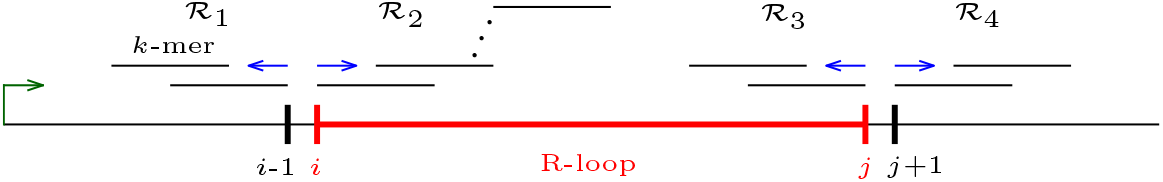
Sliding-window approach for extracting *k*-mers around the initiation and termination sites of an R-loop [*i, j*].

We tested the model for *k* = 3, 4, 5 and *p* = 7, 13. When *k* = 3, the symbol assignment in the regions r_1_ to *r*_4_ (see Figure 5 and Materials and Methods for more details) exhausts all 64 possible 3-mers and the probabilities for production rules going to *γ* or *ρ* are 0. This results in overfitting. When *k* = 5, the sequences of the two plasmids provide insufficient information, leaving between 39.1% and 58.1% of the 5-mers (*p* = 7, 13) assigned with an indeterminate symbol *γ* or *ρ*. We thus use *k* = 4 as the most suitable for training the grammar. This value of k provides information for 84.9% to 95.2% of all possible k-mers, depending on the experiment and on the choice *p* = 7 or *p* = 13. We focus on p = 13 because the choice (*k, p*) = (4, 13) provides the largest coverage for the union dictionary across all topologies (see SI Appendix Table S2).

Once the union dictionary is fixed, we use the R-loop grammar model to define a probability of R-loop formation at each nucleotide position along each plasmid. For each topology and each plasmid, first we randomly partition the data into three sets of equal size, designating two thirds of the data to define and train the grammar model. We set aside the remaining third as a *holdout set* for testing purposes. Next, we generate 30 grammar models, where we train each model on a 10% sample of the training set, taken without replacement. We then compute the probability that each nucleotide on the gene region is inside an R-loop. The final predictions are the average probabilities taken over the ensemble of 30 models (Figure 7) [24]. We tested the stability of the ensemble predictions using 3-fold cross-validation [25] (see Materials and Methods and SI Appendix Figure S6).

The R-loop grammar shows overall better prediction capabilities than the existing thermodynamics-based model R-looper [13] for both plasmids and all topologies. When compared to R-looper, our approach reduces the Root Mean Square Deviation (RMSD) by up to 55% and, in most cases it improves the Pearson correlation coefficient by at least 2-fold. Note that the thermodynamics model R-looper [13] is not trained on data. Besides the probability prediction, R-looper also provides an energy landscape for the sequence, which may need to be taken in consideration together with the probability landscape. We refer the reader to the SI Appendix for details. See Table S5 for a comparison against the holdout set and Table S6 for comparison against the full dataset. Table S7 compares the holdout set with the full dataset, and Table S8 gives the same information as Table S5 but for the deterministic symbol assignments.

The grammar rule probabilities vary depending on the plasmid topologies, and thus produce different predictions (Figure 7). Overall the fit to the data is outstanding, with Pearson correlation values from 0.68952 to 0.95165 when compared to the holdout set (Table S5). The R-loop grammar accurately identifies R-loop clusters along the gene regions in both plasmids and predicts the shift to the left as the supercoiling density increases. As noted in [26], experimental data for hypernegatively supercoiled pFC8 plasmids present with a much larger number of R-loops near the promoter region as compared with the supercoiled or linear plasmids, where they are largely absent.

## Discussion

The experimental data for hyper-negatively supercoiled pFC8 plasmids shows two clusters near the promoter region (Figure 7, bottom right). The fact that the R-loop grammar predicts one wider cluster near the promoter region can be an artifact of the model.

When processing the experimental data for training, the R-loop grammar assumed that each molecule presents with a single R-loop. On rare instances, the dataset includes molecules with more than one R-loop. One could update the grammar to include more than one R-loop. That is beyond the scope of this paper.

Plasmid topology is inherently part of the R-looper model [26]. However, the R-looper predictions for hyper-negatively supercoiled plasmids significantly underperform in detection of the early appearance of R-loops observed experimentally. Although plasmid topology is not encoded in our R-loop grammar, the model has learned the effect of supercoiling from the data and developed different probability values that distinguish between plasmid topologies.

In this work we trained the R-loop grammar on a restrictive set of plasmids and topologies, where it performs very well. However, while this is the only SMRF-seq R-loop data available to date, the plasmid sequences for pFC53 and pFC8 are not representative of the much larger set of gene sequences. As more experimental data with a larger array of genomic sequences become avalable, we anticipate that our approach will be an effective universal tool to analyze R-loop formation.

In [27] we showed that the R-loop grammar produces a set of sequences that is regular [28]. Therefore a probabilistic version of this grammar can be described by a Markov chain. This opens the door to a variety of well established techniques (e.g. [29]).

All of the code and the training data is available with complete documentation [22].

## Materials and Methods

It is known that the initiation of an R-loop is influenced by favorable G-rich DNA sequences [30, 31], while sequences spanning the lengths of R-loops may be less favorable [31, 8]. However, any patterns defining possible R-loop termination sequences are currently unknown. To identify preferable DNA sequence patterns that are specific to the initiation, elongation, and termination of R-loops, we carry out an analysis of the experimental results from [13].

### Experimental data

The sequences for pFC53 and pFC8 are reported in [13]. These plasmids share the same backbone and incorporate specific regions known to be prone to R-loop formation [32]. More specifically, pFC53 contains a 1.4-kb portion of the murine Airn CpG island, and pFC8 contains a 950-bp portion of the human SNRPN CpG island. R-loop locations along pFC53 and pFC8 are detected by SMRF-seq, a method that profiles individual R-loops at ultra-deep coverage [18, 8]. We make the complete R-loop experimental data and software available on GitHub [22].

#### *k*-mer extraction for training

Assuming that, if needed, the termination indices of all R-loops are modified so that each R-loop length is a multiple of *k* (see SI Appendix 1), we describe *k*-mers extraction around the initiation and termination sites of an R-loop.

We employ a sliding-window approach to extract the k-mers specific to the initiation, elongation, and termination of R-loops in the training set 𝒯. Let *p* ∈ ℤ^+^ be a given *padding parameter*. Consider an R-loop [*i, j*]. The regions of interest are given by *r*_1_ = [*i* − *k* − *p, i* − 1], *r*_2_ = [*i, i* + *k* + *p* − 1], *r*_3_ = [*j* − *k* − *p*+1, *j*] and *r*_4_ = [*j* +1,*j* +*k* +*p*]. We begin by taking the *k*-mer [*i*− *k*, −*i*−1] containing the k nucleotides *i* − *k*, …, *i* − 1 appearing before the beginning of the R-loop. We shift this window to the left one nucleotide at a time, for a total of p shifts. This procedure is performed while the *k*-mer remains in the gene sequence, that is, we discard any extracted *k*-mers that are not fully contained within the gene sequence. The collection of *k*-mers obtained in this way corresponds to the set of *k*-mers within region *r*_1_ and is denoted ℛ_1_. Similarly, we construct the remaining three collections of k-mers ℛ_2_, ℛ_3_, and ℛ_4_ corresponding to *k*-mers within regions *r*_2_, *r*_3_ and *r*_4_, respectively. ℛ_1_ consists of the *k*-mers preceding the beginning of the R-loops in the training set 𝒯. Similarly, ℛ_2_ (respectively, ℛ_3_ and ℛ_4_) consists of the *k*-mers at the beginning (respectively, before and after the end) of the R-loops in 𝒯. This sliding-window approach is illustrated in Figure 8.

### Selecting the most relevant *k*-mers

The *k*-mers used for training the grammar are selected from the collections ℛ_*i*_, *i* = 1, 2, 3, 4. To each *k*-mer *s* in ℛ_*i*_ we associate *weight w*_*i*_(*s*) with respect to region *r*_*i*_ as follows:

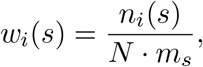

where *N* is the number of R-loops in the training set 𝒯, *n*_*i*_(*s*) is the number of occurrences of *s* in ℛ_*i*_ among all the R-loops in 𝒯, and *m*_*s*_ is the number of occurrences counted with multiplicity of *s* within the gene region of the given plasmid. This weight quantifies the prevalence of *s* in ℛ_*i*_ across all the R-loops in the training set 𝒯.

The *k*-mers in ℛ_*i*_ are ordered by weight in decreasing order *w*_*i*_(*s*_1_) > *w*_*i*_(*s*_2_) > … such that *s*_*i*_ has the *i*th highest weight. Table 1 shows an extract of all 115 4-mers in ℛ_4_ from the hyper-negatively supercoiled pFC53 R-loop data set.

**Table 1.** Selection of the most relevant 4-mers from the hyper-negatively supercoiled pFC53 R-loop data set. The first column indicates *n*, the ranking of the 4-mers in ℛ_4_ after ordering them by weight. The second column lists all the 4-mers for each *s*_*n*_; the total number of 4-mers in ℛ_4_ is 115. The third column indicates the weight *w*_4_(*s*_*n*_) of *s*_*n*_. The last three columns illustrate the steps of the selection procedure for determining the cutoff point: weight rescaling, entropy calculation, and average entropy computation. The cutoff point, highlighted with the blue background, is the maximum of the average entropy values.

| $n$ | $s_n$ | $w_4(s_n)$ | $w'_4(s_n)$ | $H_n$ | $h_n$ |
| --- | --- | --- | --- | --- | --- |
| 1 | CAAT | 0.03571 | 1.00000 | 0.00000 | 0.00000 |
| 2 | GGAT | 0.03214 | 0.90000 | 0.04118 | 0.02059 |
| 3 | AGGT | 0.02976 | 0.83333 | 0.06598 | 0.03572 |
| ⋮ | | | | | |
| 29 | GAAG, CCGT,<br>CGCA, ACCA | 0.00893 | 0.25000 | 0.15051 | 0.14094 |
| 30 | CAAG, AAGC, GGTT | 0.00824 | 0.23077 | 0.14696 | 0.14147 |
| 31 | CCCG | 0.00794 | 0.22222 | 0.14516 | 0.14168 |
| 32 | CTCT, TTCA, CGGA,<br>ATTT, TACA, GTGC,<br>GAGT, GCGT,<br>CGTG, GGTA, GTAG,<br>GCCA, AGCC | 0.00714 | 0.20000 | 0.13979 | 0.14165 |
| 33 | AAGG | 0.00630 | 0.17647 | 0.13294 | 0.14128 |
| ⋮ | | | | | |
| 40 | GCCC | 0.00298 | 0.08333 | 0.08993 | 0.13399 |
| 41 | AAAG | 0.00275 | 0.07692 | 0.08569 | 0.13357 |
| 42 | GCAC | 0.00255 | 0.07143 | 0.08187 | 0.13312 |

Total number of 4-mers = 115

In order to identify the most relevant *k*-mers in each ℛ_*i*_, we determine a cutoff point for thresholding the ordered list ℛ_*i*_ using a procedure that relies on entropy reduction [33]. We rescale all weights of the *k*-mers in ℛ_*i*_ by normalizing with respect to the heighest weight (i.e. *s*_1_) with 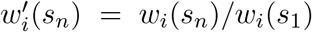. We consider the entropy of the *s*_*n*_ as 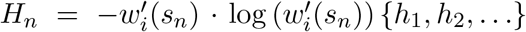 and compute the average entropy of *s*_1_,…, *s*_*n*_ as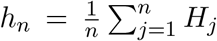. The (global) maximum of {*h*_1_, *h*_2_,…} [33] * is set to be the threshold of ℛ_*i*_. The threshold reduced list of k-mers 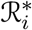, called *highly weighted*, comprises of all k-mers *s*_𝓁_ in ℛ_*i*_ such that *h*_𝓁_ is greater than or equal to the threshold value of ℛ_*i*_^†^.

### Training set dictionary

For each plasmid and each set of experimental conditions, we use the list of *k*-mers together with their associated weights to form a dictionary that enables us to represent each R-loop with an R-loop grammar word.

Given an R-loop [*i, j*] within the gene region [*b, e*], we focus on three segments [*b, i* −1], [*i, j*], and [*j* + 1, *e*] which comprise the sequences preceding, within, or following the R-loop respectively. We subdivide each of the three nucleotide sequences into consecutive and non-overlapping *parsing blocks*. These blocks are *k*-mers, except possibly for the block that ends with *i* − 1 and the one that starts with *j* + 1, which could be shorter (i.e., of length mod k). By construction (SI Appendix Section 1) the R-loop segment [*i*, j] is always a multiple of *k*. See Figure 9.

**Figure 9.**
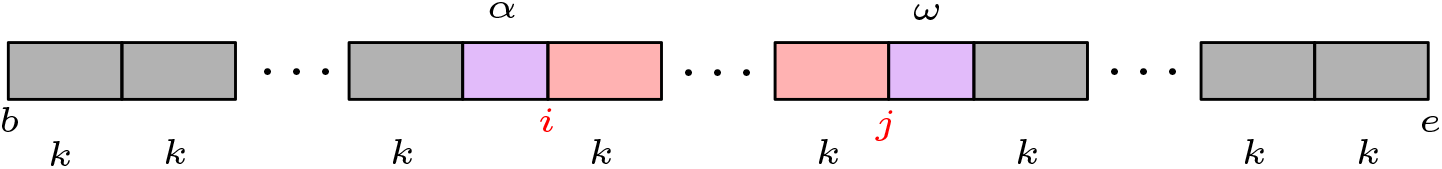
Subdivision of the gene region into parsing blocks; *i* and *j* indicate the initiation and the termination of the R-loop, respectively.

To write a nucleotide sequence containing an R-loop as a word in the R-loop grammar, we determine correspondence between the parsing blocks within [*b, i* − 1], [*i, j*], and [*j* + 1, *e*] and grammar symbols. This correspondence between parsing blocks and grammar symbols, called a *dictionary* 𝒟, is obtained through a symbol assignment function *C*(𝓁, *s*) depending on the location 𝓁 of the first nucleotide of the *k*-mer s.

The grammar symbol assignment depends on the parsing block weights generated by the training set 𝒯. For a given training set 𝒯, a parsing block can be highly weighted in some regions (*r*_*i*_,*i* = 1, 2, 3, 4, Figure 5), appearing in some regions but not-highly weighted in any of them, or not appearing in any of the regions. For example, highly weighted parsing blocks in region *r*_4_ are treated as stable DNA:DNA duplexes, and those within *r*_2_ are treated as stable RNA:DNA duplexes; however, the weighted values can imply ambiguous assignments, so we use more complex symbol assignment maps. We define the symbol assignment map *C*(𝓁, *s*) below.

#### Highly weighted parsing block assignments

Let *s* be a *k*-mer which belongs to one of the threshold reduced lists 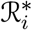. We use 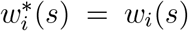 if 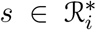 and 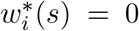 if 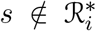. Let 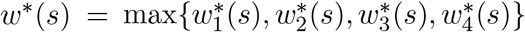. We define the map *C*(𝓁, *s*) in the following manner.

- If 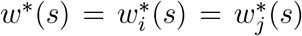 with *i* ∈ {1, 2} and *j* ∈ {3, 4} then *s* is highly weighted as it appears at both the start and the end of an R-loop. Then

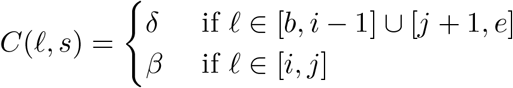

treating *s* as both a stable and an unstable DNA:DNA (RNA:DNA) hybrid when the *k*-mer is outside (resp. inside) the R-loop. Otherwise,
- If 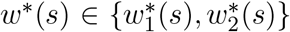 then *C*(𝓁, *s*) = *σ* for 𝓁 ∈ [*b, i* − 1] [*j* + 1, *e*] and 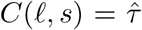 for 𝓁 ∈ [*i, j*]. This treats *k*-mers at the beginning of the R-loop as stable RNA:DNA hybrids and unstable DNA:DNA duplexes.
- If 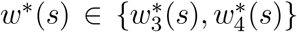 then 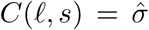 for 𝓁 ∈ [*b, i* − 1] ∪ [*j* + 1, *e*] and *C*(𝓁, *s*) = *τ* for 𝓁 ∈ [*i, j*]. Highly weighted *k*-mers at the end of the R-loop are treated as unstable RNA:DNA hybrids and stable DNA:DNA duplexes.

#### Not highly weighted parsing block assignment

Consider a *k*-mer *s* which does not appear in any of the threshold reduced lists 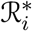. We define *w*_*i*_(*s*) = 0 if *s* ∉ ℛ_*i*_. Let *N*_*b*_, *N*_*r*_ and *N*_*e*_ be the number of times s appears as a parsing block before, within and after (resp.) an R-loop in the training set 𝒯. Let *M* = max{*N*_*b*_, *N*_*r*_, *N*_*e*_}.

- If the maximum *M* is achieved in a non-unique way, consider the maximal weight *w*(*s*) = max *w*_1_(*s*), *w*_2_(s), *w*_3_(s), *w*_4_(s). If *w*(*s*) = 0, set *C*(𝓁, *s*) = *γ* for 𝓁 ∈ [*b, i* − 1] ∪ [*j* + 1, *e*] or *C*(𝓁, *s*) = *ρ* for 𝓁 ∈ [*i, j*] implying that we cannot determine whether this *k*-mer is (un)stable DNA:DNA duplex, or RNA:DNA hybrid (resp.). If *w*(*s*) ≠ 0 we proceed according to the weights as in the case of highly weighted k-mers.

Otherwise, *s* has a maximal number of appearances in a unique region. Then we treat the number of occurrences of *s* within a region as weights, in particular,

- if *M* = *N*_*b*_ then *C*(𝓁, *s*) = *σ* for 𝓁 ∈ [*b, i* - 1] ∪ [*j* + 1, *e*] and 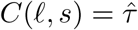 for 𝓁 ∈ [*i, j*];
- if *M* = *N*_*e*_ then 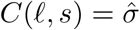 for 𝓁 ∈ [*b, i* − 1] ∪ [*j* + 1, *e*] and *C*(𝓁, *s*) = *τ* for 𝓁 ∈ [*i, j*];
- if *M* = *N*_*r*_ we focus on the weights *w*_2_(*s*), *w*_3_(*s*) in regions *r*_2_, *r*_3_ resp. and pursue as in the case of highly weighted *k*-mers.

### Union of dictionaries

For the experimental data obtained by *in vitro* transcription of both plasmids pFC53 and pFC8 and the detected R-loops for the same topological state we obtain two dictionaries 𝒟_1_ and 𝒟_2_. We form a *union dictionary*, 𝒟, which incorporates the information from both 𝒟_1_ and 𝒟_2_. Let *C*(𝓁, *s*) (respectively, *C*^(1)^(𝓁, *s*), *C*^(2)^(𝓁, *s*)) be the symbol assignments for 𝒟 (respectively, 𝒟_1_, 𝒟_2_). The symbol assignment *C* for the dictionary 𝒟 is based on the symbol assignments by *C*^(1)^ and *C*^(2)^. If the symbol assignment of both *C*^(1)^(𝓁, *s*) and *C*^(2)^(𝓁, *s*) is the same, we assign the same symbol *C*(𝓁, *s*). When the symbol assignment by *C*^(1)^(𝓁, *s*) and *C*^(2)^(𝓁, *s*) differ, we apply two approaches to resolve the conflict - a deterministic one and a stochastic one (SI Appendix 2).

### R-loops as words using a dictionary

For training the grammar, we write all experimental R-loops as words over the alphabet 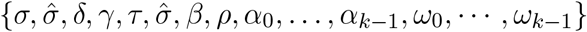. For producing the model predictions, we write all possible R-loops as words over the same alphabet. This is done by first splitting the gene into parsing blocks as in Figure 9 according to the initiation and termination indices [*i, j*]. All but two of the blocks are *k*-mers and in those cases the symbol is assigned according to the appropriate section of the dictionary, depending on whether the block is before, within, or after the positions [*i, j*]. The lengths of the blocks *α* and *ω* for the transitions can vary between 0 and *k* − 1 depending on the values *i,j,k*, so we assign a symbol *α* _*l*_ or *ω*_*l*_, respectively, to indicate the length of the block. Note that this means that each R-loop word contains exactly one *α* and *ω* even when the corresponding length of the gene sequence is 0.

## Acknowledgments

The whole team acknowledges support from the National Science Foundation and the National Institutes of Health, DMS/NIGMS awards #2054347 and #2054321. In addition, MV acknowledges support from NSF grants DMS#1716987 and DMS#1817156, SP acknowledges support by Simons Foundation gift MP-TSM-00002798 and NSF DMS#1815832, MMF acknowledges the support of the Natural Sciences and Engineering Research Council of Canada (NSERC) [funding reference numbers DGECR-2023-00131, RGPIN-2023-04722] and the University of Manitoba (research start-up funds) and NJ acknowledges support in part by NSF CCF#2107267, the W.M. Keck Foundation and the Center for Mathematics of Complex Biological Systems under NSF grant DMS-1764406 and Simons Foundation Grant 594594. FC and SH were also supported in part by NIH grant R35 GM139549. MMF, NJ, SP, MR and MV thank the Institute of Pure and Applied Mathematics and the Association for Women in Mathematics for seeding this research.

## Supporting Information

### 1. R-loop data pre-processing

We provide all data files and data extraction details in the GitHub repository (1). The template strand 5′ − 3′ of each plasmid is in FASTA format and the corresponding R-loop locations for each of the three topologies considered are included in BED files. In the following exposition we consider each plasmid in the 5′ −3′ direction of the non-template strand while R-loop locations are specified by their initiation and termination indices, *i* and *j*, with *j > i*. We denote the R-loop segment as interval of nucleotides [*i, j*], i.e., the sequence of nucleotides *i*, …, *j*.

We use *k*-mers (*k* ∈ ℤ^+^) to analyze sequence preference for initiation, elongation, and termination of R-loops and assume that the R-loops can be parsed with *k*-mers. So for each R-loop [*i, j*], we increase or decrease (whichever modification is smaller) the termination index *j* to make the R-loop length a multiple of *k*. For even values of *k* if there are two equal modifications of the termination, we choose to slightly elongate the R-loop. Regardless of the parity of *k*, if the elongation falls outside of the gene region (past the transcription end) then we choose the new termination index to decrease the R-loop size. Figure S1 shows a schematic representation of this pre-processing of R-loop windows.

**Fig. S1.**
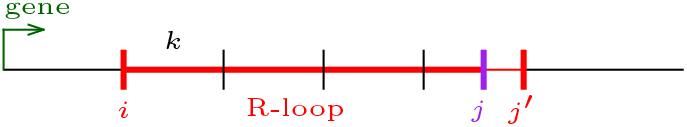
Modification of an R-loop termination index. The experimentally detected R-loop, red thick line, spans the sequence [*i, j*]. Here the new termination index *j′* increases the termination index *j* such that the resulting R-loop length is a multiple of *k* spanning sequence [*i, j′*]. Black horizontal line: DNA non-template strand of the plasmid; green arrow: transcription direction.

### 2. R-loop grammar symbol assignments

#### A. Defining the Dictionaries

The *General approach* and the Materials and Methods sections in the main text give an overview of the process for defining the dictionaries. Here we give more details.

The dictionaries obtained after training on each plasmid have a proportion of indeterminate symbols ranging from 15% to 30.2% (see Table S1). This results in poor prediction value when training on one plasmid and testing on the other. To improve the prediction we incorporate the information from both plasmids by taking a union of dictionaries. The union of dictionaries have a proportion of indeterminate symbols ranging from 4.8% to 13.4% (Table S1). When taking the union, we consider two ways of resolving conflicting information in symbol assignment, stochastic (coin toss) and deterministic, guided by the description of the grammar symbols.

We determine the symbol assignment in the union of dictionaries as follows. If *C*^(1)^(𝓁, *s*)= *C*^(2)^(𝓁, *s*), we set *C*(𝓁, *s*)= *C*^(1)^(𝓁, *s*). If the symbol assignment conflicts between *C*^(1)^ and *C*^(2)^, i.e. *C*^(1)^(𝓁, *s*) ≠ *C*^(2)^(𝓁, *s*), the assignments are resolved by examining the weights of the *k*-mer *s* where the two dictionaries disagree, with one set of weights computed from the pFC53 (*seq*_1_) experimental data and the other from pFC8 (*seq*_2_) data. We use one of the deterministic or stochastic ways to resolve the conflicts. On average, in our computations there were 6-8 *k*-mers with conflicting symbol assignments.

##### A.1 Deterministic union of dictionaries

We resolve conflicts between *C*^(1)^(𝓁, *s*) and *C*^(2)^(𝓁, *s*) based on the meaning of the grammar symbols as explained in section *Formal grammars and R-loops* in the main text.

More precisely, for a *k*-mer *s* and 𝓁 ∈ [*b, i* − 1] ∪ [*j* + 1, *e*],

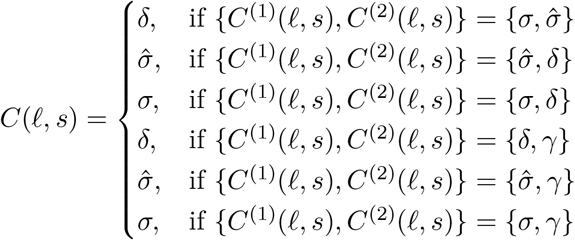

We define the assignment to *k*-mers inside the R-loop, when 𝓁 ∈ [*i, j*], by

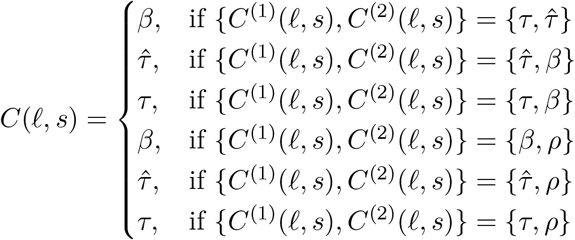

##### A.2. Stochastic union of dictionaries

When there is not enough information we perform a random coin toss to resolve the conflict. In the case where *w*^(1)^(*s*) = *w*^(2)^(*s*), we randomly set *C*(*𝓁, s*) to be *C*^(1)^(*𝓁, s*) or *C*^(2)^(∈, *s*).

### 3. Training the grammar

#### A. Training on one sequence

We associate a probability to each production rule in the grammar, thereby creating a stochastic grammar. The probabilities of the rules with the same non-terminal sum up to 1.

We determine the rule probabilities by the frequency of each rule used in the derivations of the words that correspond to R-loops from the training set (10% chosen randomly from 2*β* of the data for each plasmid). The grammar is non-ambiguous, i.e. each R-loop word has precisely one parse. Therefore, given a word in the grammar one can easily reverse engineer the sequence of rules that generate the word.

Given a training set of *N* R-loops, let {*v*_1_, *v*_2_,…, *v*_*N*_} be the set of its corresponding words. By construction, each of them is of the form

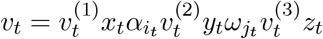

where 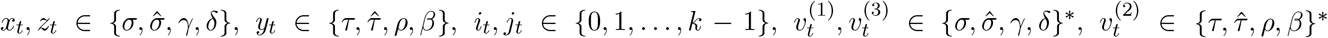. The production rules starting with the starting non-terminal *S* are used to derive the segments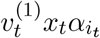.

For a word *v*, we use the standard notation |*v*| to indicate the length of *v* and |*v*|_*a*_ to indicate the number of symbols *a* appearing in *v*. For each non-terminal *X* we denote with *q*(*X*) the number of different types of rules of the form *X* → *r*, where *r* is any word consisting of terminals and non-terminals. So, *q*(*S*)= *q*(*R*)= *q*(*Q*)= 8. We use Laplace smoothing (2) to avoid cases with probability 0 (in those cases we take *η* = 1). We set

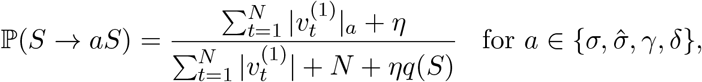

and

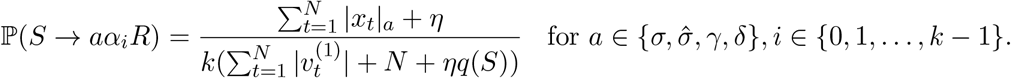

We derive the R-loop via productions starting with the non-terminal *R*, which yield the segments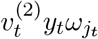. Therefore, we set

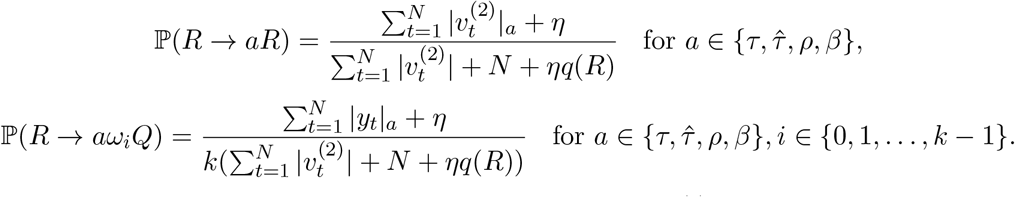

The non-terminal *Q* is used to derive *z*_*t*_ and the remaining string 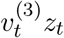. So we set

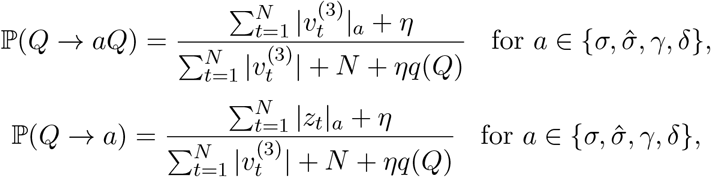

#### B. Training on two sequences

Suppose 𝒯_1_ is a training set with *N*_1_ R-loops for sequence *seq*_1_ and 𝒯_2_ is a training set with *N*_2_ loops for sequence *seq*_2_. We constructed dictionaries 𝒟_1_ and 𝒟_2_ and let dictionary 𝒟 be the union of 𝒟_1_ and 𝒟_2_, as described in the main text (Materials and Methods). For each of the rules *A* → *w* of the grammar (see section *Symbol assignment and R-loop production rules* in the main text), where *A* is a non-terminal symbol and *w* is a word, we compute two probabilities, 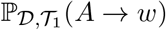 and 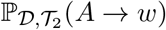 as above. Finally, we set

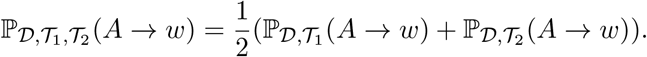

### 4. Model predictions

We divide the data into thirds, and use 2/3 for training of the grammar and assignment of probabilities. We tested the prediction against the remaining 1/3 of the data (the *holdout set*). We created an ensemble of 30 different grammar models; each of them trained as described in the main text (Results). Fig. 7 shows the average probabilities predicted by the 30 runs of the grammar against the holdout set. Fig. S4 and Fig. S5 show the results of 30 runs of the grammar against the full set of data with stochastic and deterministic symbol assignments (resp.).

To test the stability of the predictions from the ensemble, we used 3-fold cross-validation. In addition to the original holdout set (fold 1), we created other two non-overlapping folds from the 2/3 of the data used for model training. Subsequently, we retrained the model using the remaining data. In each of the 3 cases, the training set comprised two-thirds of the data, while the remaining one-third constituted the test set (fold). Fig. S6 shows the predictions from each of the 3 ensembles.

---

**Fig. S2.**
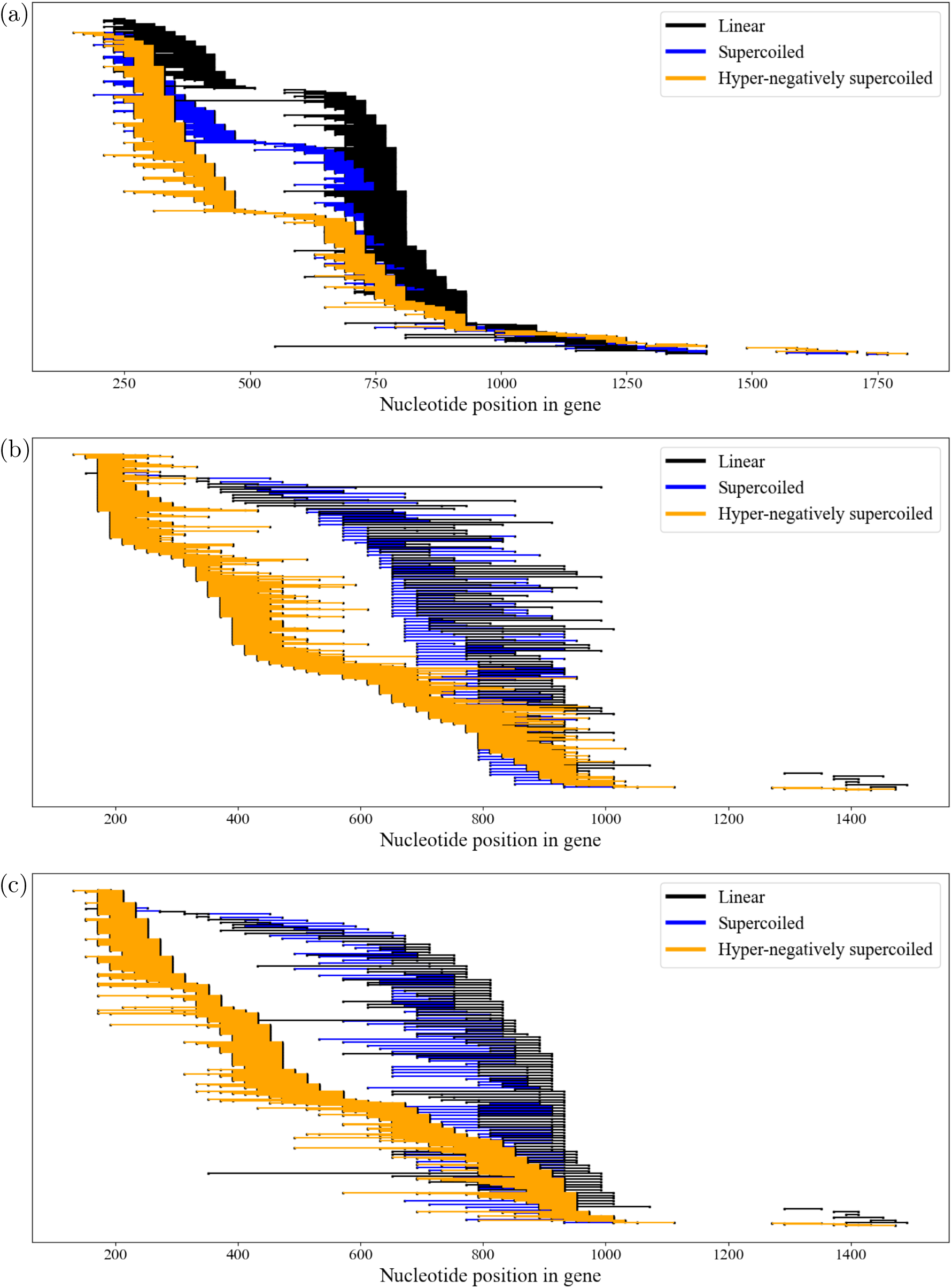
Experimental R-loop locations for plasmids pFC53 (a) and pFC8 ((b) and (c)) with starting topology: linear (black); supercoiled (blue); and hyper-negatively supercoiled (orange). The *x*-axis indicates the nucleotide position of the gene starting at 0, rounded to the nearest 20th nucleotide. Each horizontal line segment corresponds to one experimentally detected R-loop. The R-loops have been sorted by the starting (b) or ending nucleotide ((a) and (c)). Each data set is uniformly spread vertically (116 linear, 104 supercoiled and 1044 hyper-negatively supercoiled for pFC8, and 255 linear, 612 supercoiled and 408 hyper-negatively supercoiled for pFC53). Proportional differences in R-loop initiation under the three conditions can be observed independent of the number of experimental R-loops observed.

**Fig. S3.**
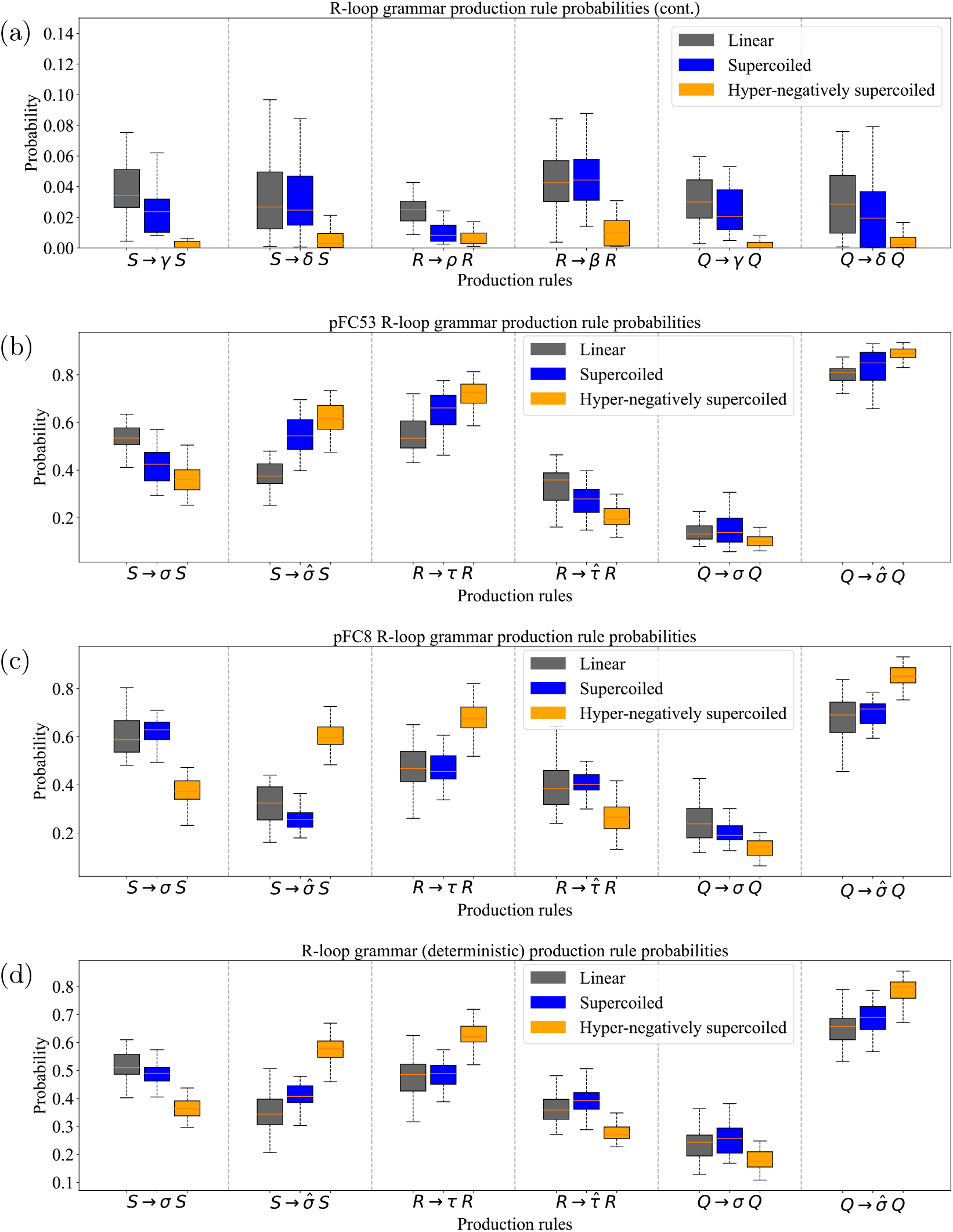
Production rule probabilities. The boxplots illustrate the changes in the probabilities for the six indeterminate production rules (a) and for the main production rules (b-d) related to the stability of the structure before, within, and after an R-loop as the topology from the substrate changes from linear to hyper-negatively supercoiled. In each case, we used the grammar defined for the union or for one of the plasmid’s dictionary with parameters *n* = 4 and *p* = 13. The mid-line of each box is the median, with the first and third quartiles indicated by the box frames. The whiskers represent the largest point not more than 1.5 interquartile range (IQR) beyond the box frame. (a) probabilities with the dictionary for pFC8 ⋃ pFC53 for the indeterminate rules not shown in Fig. 6. (b) probabilities for dictionary of pFC53 (c) probabilities for dictionary of pFC8. (d) probabilities for the deterministic union (pFC8 ⋃ pFC53).

**Fig. S4.**
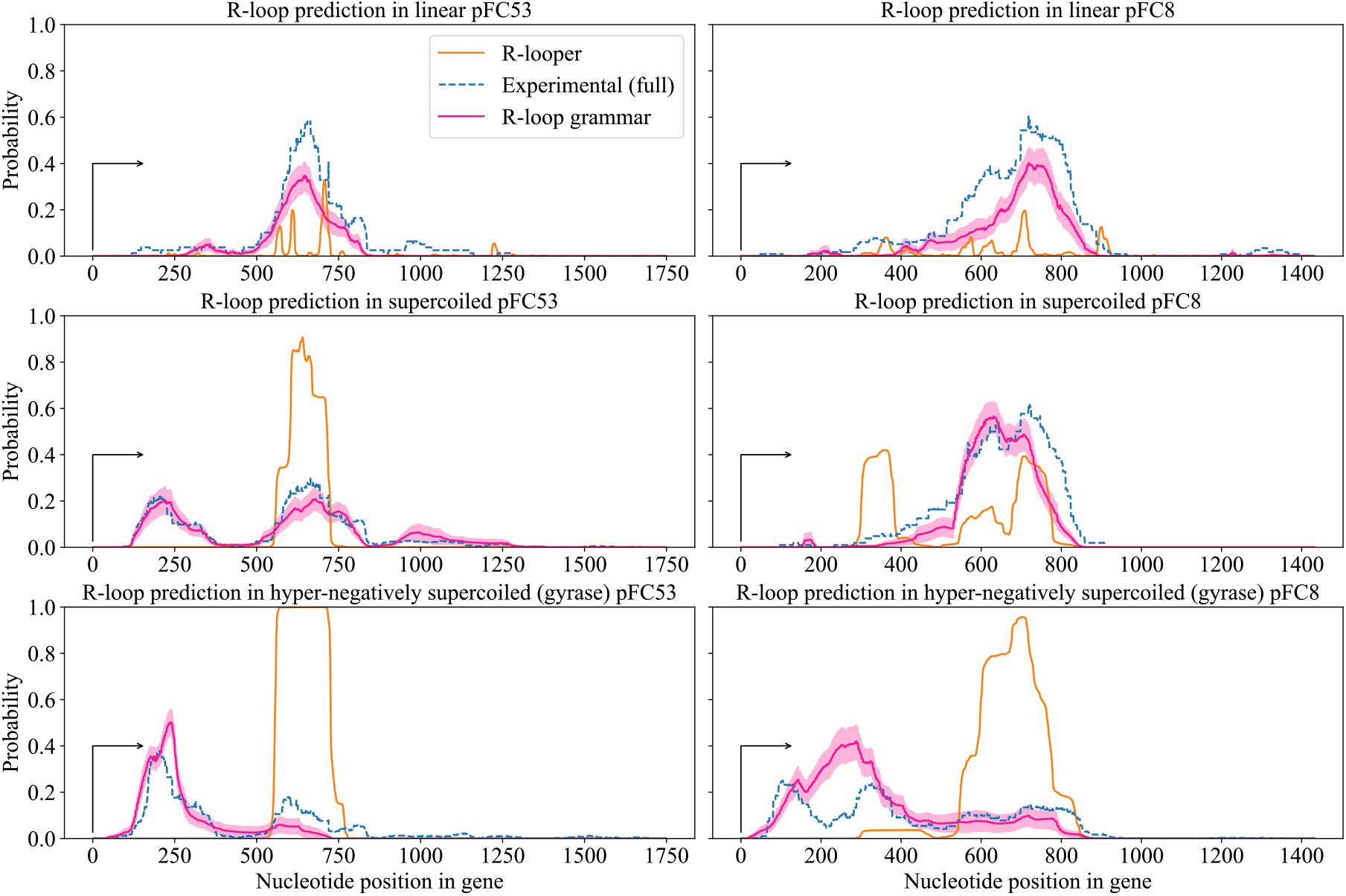
Predictions from the <u>stochastic</u> R-loop grammar model and from R-looper for different topologies on plasmids pFC8 and pFC53 against the full experimental dataset. The graphs show the predictions from R-looper (orange) and the predictions from the R-loop grammar ensemble of 30 models (pink). The pink shaded area corresponds to the standard error of the mean (s.e.m.) for the ensemble. The dashed blue line shows the observed proportion of R-loops in the full experimental dataset. We indicate the substrate topology in each graph: linear (top row); supercoiled (middle row); hyper-negatively supercoiled (bottom row).

**Fig. S5.**
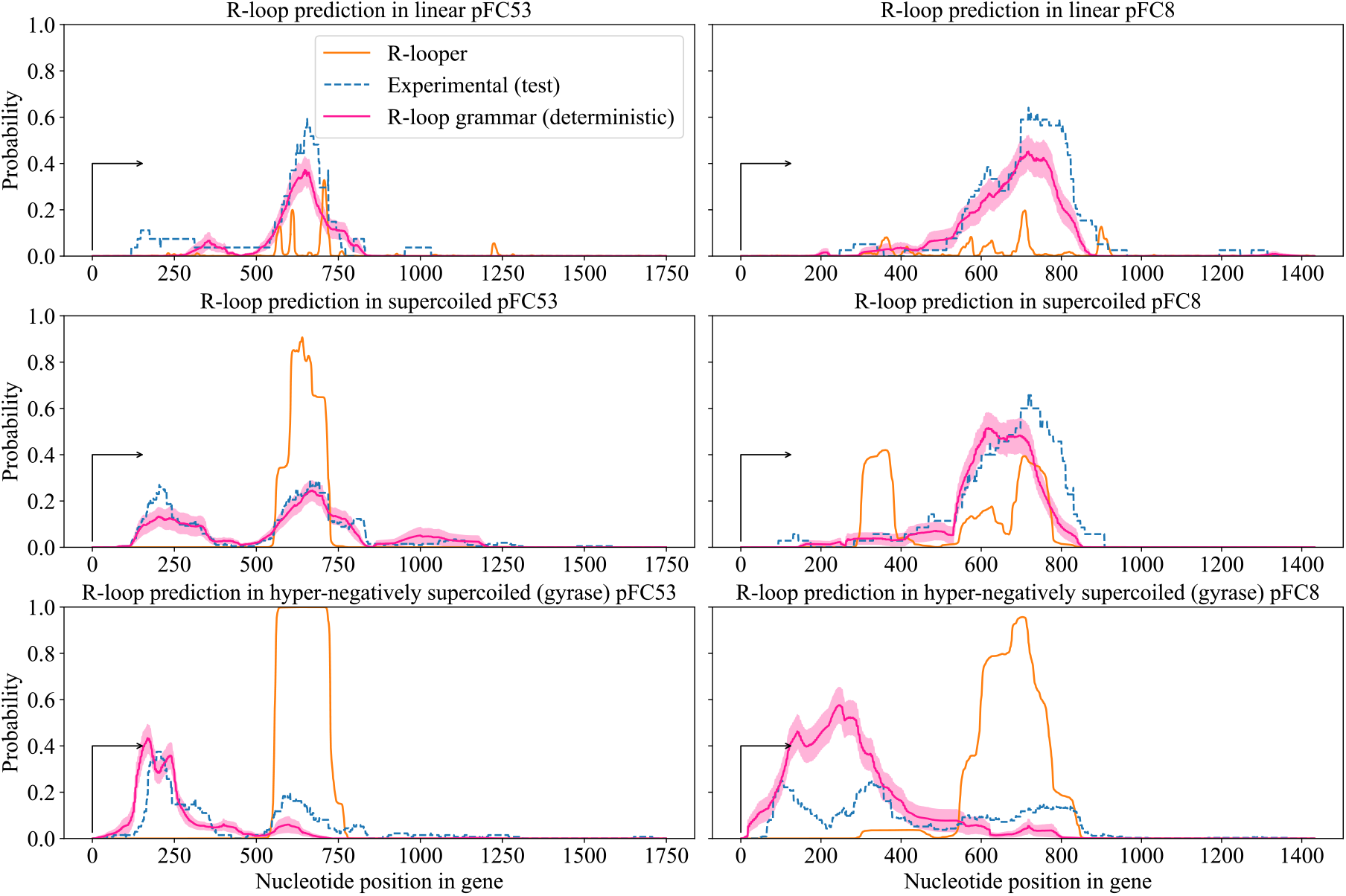
Predictions from the <u>deterministic</u> R-loop grammar model and from R-looper for different topologies on plasmids pFC8 and pFC53 against the holdout set. The graphs show the predictions from R-looper (orange) and the predictions from the R-loop grammar ensemble of 30 models (pink). The pink shaded area corresponds to the s.e.m. for the ensemble. The dashed blue line shows the observed proportion of R-loops in the holdout set. We indicate the substrate topology in each graph: linear (top row); supercoiled (middle row); hyper-negatively supercoiled (bottom row).

**Fig. S6.**
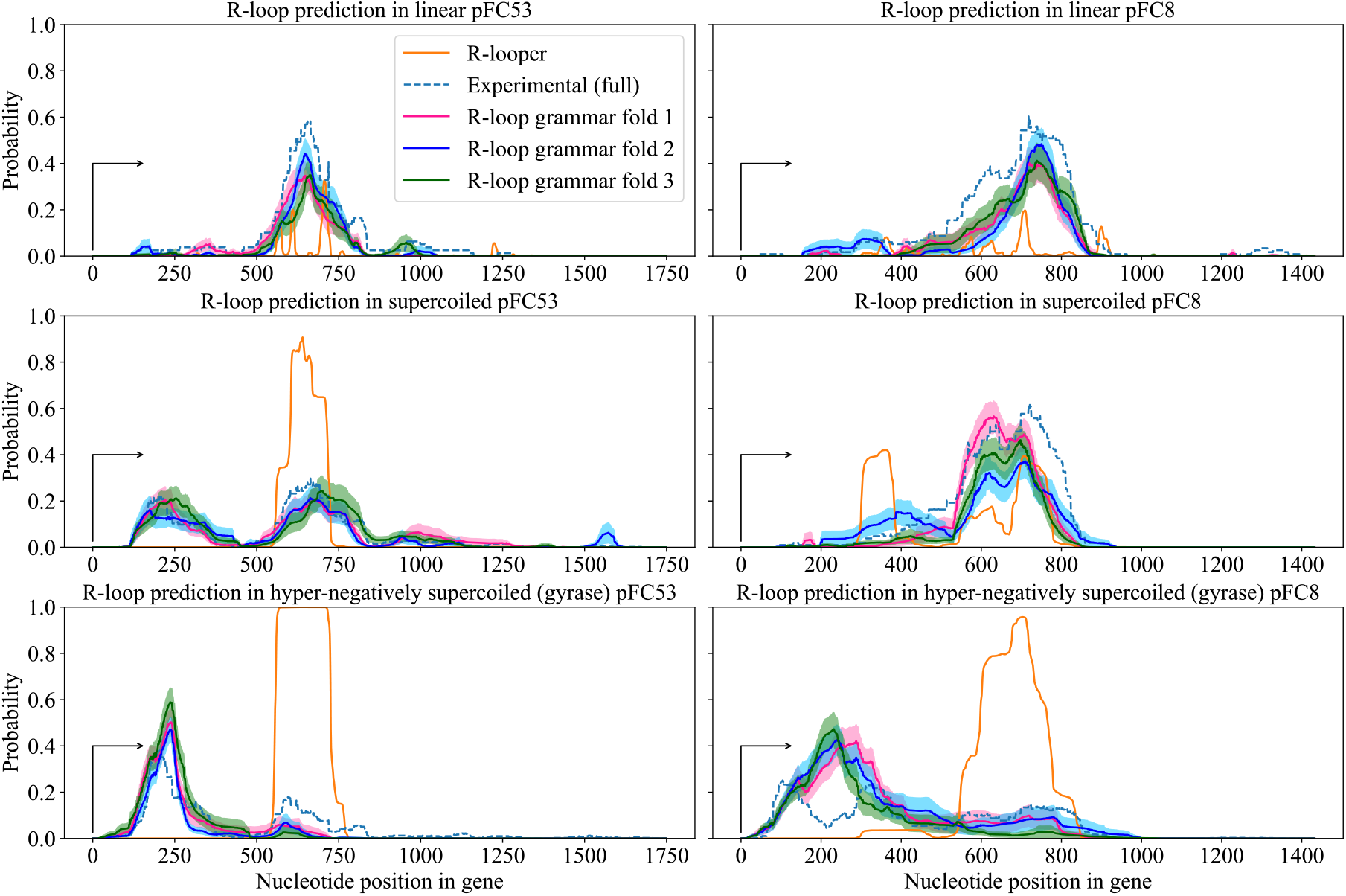
3-fold predictions from the <u>stochastic</u> R-loop grammar model trained on three distinct two thirds of the data. We computed predictions for different topologies on plasmids pFC8 and pFC53. The graphs show the predictions from each fold from the R-loop grammar ensemble of 30 models (pink, blue, green). The shaded areas correspond to the s.e.m. for each fold ensemble. The orange line corresponds to predictions from R-looper. The dashed blue line shows the observed proportion of R-loops in the full experimental dataset. We indicate the substrate topology in each graph: linear (top row); supercoiled (middle row); hyper-negatively supercoiled (bottom row).

**Table S1.** Average *k*-mer coverage for determinate symbols (i.e. not *γ* nor *ρ*) calculated by looking at the individual dictionaries obtained when training the R-loop grammar (with *k* =4 and *p* = 13) on data from the plasmids pFC8 (row 1), pFC53 (row 2), and when taking the union of the dictionaries. For each dictionary, we focused on *k*-mers that are assigned a symbol that is not one of the indeterminate symbols. There are a total possible 44 = 256 assignments. We computed the average assignment coverage over the ensemble of 30 runs. The results are reported with the Mean% (SD), where SD is the standard deviation.

| Training on | Topology |  |  |
| --- | --- | --- | --- |
|  | Linear | Supercoiled | Hyper-negatively supercoiled |
| pFC8 | 72.1% (2.4) | 69.8% (1.6) | 85.0% (1.2) |
| pFC53 | 72.4% (2.0) | 82.7% (1.9) | 83.0% (2.5) |
| pFC8 ∪ pFC53 | 86.6% (1.5) | 91.4% (1.3) | 95.2% (0.9) |

**Table S2.** Average *k*-mer coverage for determinate symbols calculated for the union of dictionaries for different pairs of parameters (*k, p*). We look at instances where *k*-mers are assigned a symbol that is not *γ* or *ρ* in the union of dictionaries. We tested the model for *k* = 3, 4, 5 and *p* = 7, 13. When *k* =3 the symbol assignment exhausts all 64 possible 3-mers and the probabilities for production rules going to *γ* or *ρ* are 0 (data not shown). This results in overfitting. Here we show the average determinate coverage for *k* = 4, 5 and *p* = 7, 13. There are a total possible 44 = 256 assignments for *k* =4 and 45 = 1024 assignments for *k* = 5, and we look at the average assignment coverage over the ensemble of 30 runs. The results are reported with Mean% (SD), where SD is the standard deviation. The choice (*k, p*)= (4, 13) provides the largest coverage for the union dictionary across all topologies.

| Topology | $k = 4$ | | $k = 5$ | |
| --- | --- | --- | --- | --- |
| | $p = 7$ | $p = 13$ | $p = 7$ | $p = 13$ |
| Linear | 84.9% (3.7) | 86.6% (1.5) | 41.9% (3.3) | 43.3% (3.3) |
| Supercoiled | 90.2% (3.8) | 91.4% (1.3) | 48.9% (1.4) | 50.2% (1.3) |
| Hyper-negatively supercoiled | 94.2% (3.0) | 95.2% (0.9) | 59.1% (1.6) | 60.9% (1.8) |

**Table S3.** Statistical comparison of production rule probabilities across topologies. After obtaining significant ANOVA results for each production rule, we conducted pairwise *t*-tests across all topology pairs. We adjusted the p-values for multiple comparisons using the Bonferroni method.

| Production Rule | Linear vs. supercoiled | Linear vs. hyper-negatively supercoiled | Supercoiled vs. hyper-negatively supercoiled |
| --- | --- | --- | --- |
| $S \rightarrow \sigma S$ | 0.0788 | 0.0000 | 0.0000 |
| $S \rightarrow \hat{\sigma} S$ | 0.0060 | 0.0000 | 0.0000 |
| $R \rightarrow \tau R$ | 1.0000 | 0.0000 | 0.0000 |
| $R \rightarrow \hat{\tau} R$ | 0.6309 | 0.0000 | 0.0000 |
| $Q \rightarrow \sigma Q$ | 1.0000 | 0.0000 | 0.0001 |
| $Q \rightarrow \hat{\sigma} Q$ | 0.7796 | 0.0000 | 0.0000 |

**Table S4.** Spearman and Kendall’s Tau correlations for each production rule. The correlation is computed between the rule probabilities values and the corresponding topologies according to the level of supercoiling.

| Production Rule | Spearman | p-value | Kendall's Tau | p-value |
| --- | --- | --- | --- | --- |
| $S \rightarrow \sigma S$ | -0.75 | 0.0000 | -0.61 | 0.0000 |
| $S \rightarrow \hat{\sigma} S$ | 0.80 | 0.0000 | 0.66 | 0.0000 |
| $R \rightarrow \tau R$ | 0.67 | 0.0000 | 0.52 | 0.0000 |
| $R \rightarrow \hat{\tau} R$ | -0.53 | 0.0000 | -0.40 | 0.0000 |
| $Q \rightarrow \sigma Q$ | -0.48 | 0.0000 | -0.37 | 0.0000 |
| $Q \rightarrow \hat{\sigma} Q$ | 0.65 | 0.0000 | 0.51 | 0.0000 |

**Table S5.**
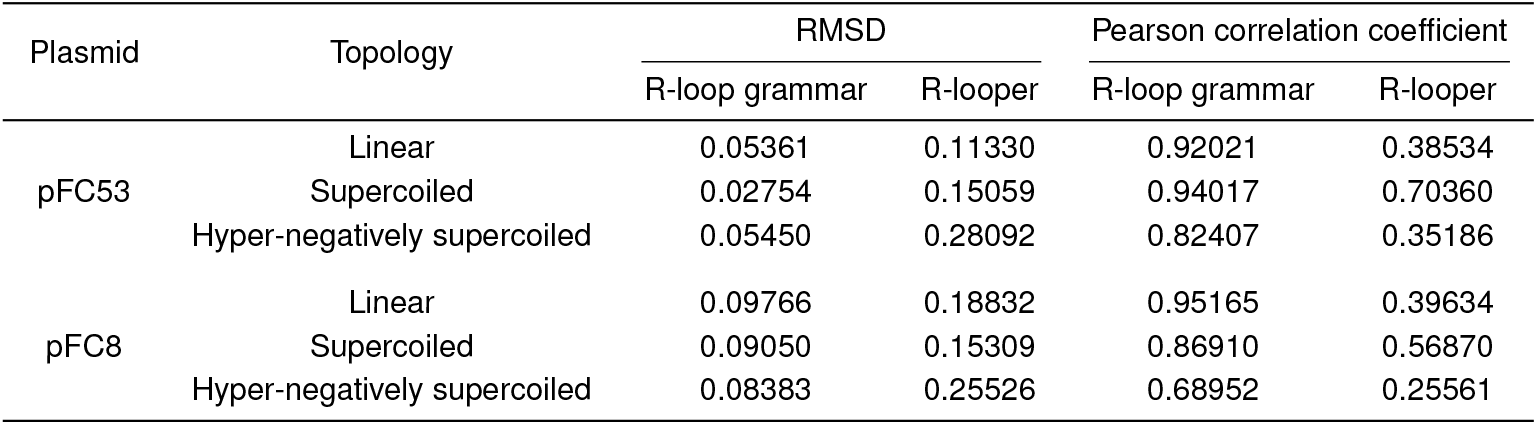
RMSD and Pearson correlation coefficient calculated by comparing the predictions obtained using the R-loop grammar (union of dictionaries; parameters *k* =4 and *p* = 13) and R-looper <u>against the holdout set</u>.

**Table S6.** RMSD and Pearson correlation coefficient calculated by comparing the predictions obtained using the R-loop grammar (union of dictionaries; parameters *k* =4 and *p* = 13) and R-looper <u>against the full set</u>.

| Plasmid | Topology | RMSD |  | Pearson correlation coefficient |  |
| --- | --- | --- | --- | --- | --- |
|  |  | R-loop grammar | R-looper | R-loop grammar | R-looper |
| pFC53 | Linear | 0.05935 | 0.12811 | 0.97337 | 0.40471 |
|  | Supercoiled | 0.02692 | 0.14704 | 0.94135 | 0.74356 |
|  | Hyper-negatively supercoiled | 0.04861 | 0.28492 | 0.86632 | 0.30082 |
| pFC8 | Linear | 0.08791 | 0.17794 | 0.95792 | 0.41878 |
|  | Supercoiled | 0.07429 | 0.15596 | 0.92038 | 0.55947 |
|  | Hyper-negatively supercoiled | 0.08544 | 0.25219 | 0.66803 | 0.29600 |

**Table S7.** Pearson correlation coefficient calculated by comparing the predictions obtained using the R-loop grammar (union of dictionaries; parameters *k* =4 and *p* = 13) <u>against the holdout (test) set and the full set (full)</u>.

| Plasmid | Topology | Pearson correlation coefficient |  |
| --- | --- | --- | --- |
|  |  | R-loop grammar (test) | R-loop grammar (full) |
| pFC53 | Linear | 0.92021 | 0.97337 |
|  | Supercoiled | 0.94017 | 0.94135 |
|  | Hyper-negatively supercoiled | 0.82407 | 0.86632 |
| pFC8 | Linear | 0.95165 | 0.95792 |
|  | Supercoiled | 0.86909 | 0.92038 |
|  | Hyper-negatively supercoiled | 0.68952 | 0.66803 |

**Table S8.** RMSD and Pearson correlation coefficient calculated by comparing the predictions obtained using the <u>deterministic</u> symbol assignments for the R-loop grammar (union of dictionaries; parameters *k* =4 and *p* = 13) and R-looper <u>against the the holdout set</u>.

| Plasmid | Topology | RMSD |  | Pearson correlation coefficient |  |
| --- | --- | --- | --- | --- | --- |
|  |  | R-loop grammar (deterministic) | R-looper | R-loop grammar (deterministic) | R-looper |
| pFC53 | Linear | 0.05204 | 0.11331 | 0.93690 | 0.38534 |
|  | Supercoiled | 0.02876 | 0.15059 | 0.93565 | 0.70360 |
|  | Hyper-negatively supercoiled | 0.05959 | 0.28092 | 0.74841 | 0.35187 |
| pFC8 | Linear | 0.07718 | 0.18832 | 0.94558 | 0.39634 |
|  | Supercoiled | 0.08669 | 0.15309 | 0.88208 | 0.56871 |
|  | Hyper-negatively supercoiled | 0.13892 | 0.25526 | 0.61379 | 0.25561 |

## Footnotes

* It is achieved when (*n* − 1)*h*_*n* ≤_ *h*_1_ + *·· ·* + *h*_*n*_ − _1_, that is, when adding a new value *h*_*n*_ is ‘not significant’ with respect to the sum of the already added values.

† There may be several *k*-mers corresponding to the threshold cutoff. In this case, all such *k*-mers are included in the highly weighted list 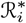.

## Notes

### Competing Interest Statement

The authors have declared no competing interest.

